# Impaired competence in flagellar mutants of *Bacillus subtilis* is connected to the regulatory network governed by DegU

**DOI:** 10.1101/194027

**Authors:** Theresa Hölscher, Tina Schiklang, Anna Dragoš, Anne-Kathrin Dietel, Christian Kost, Ákos T. Kovács

## Abstract

The competent state is a developmentally distinct phase, in which bacteria are able to take up and integrate exogenous DNA into their genome. *Bacillus subtilis* is one of the naturally competent bacterial species and the domesticated laboratory strain 168 is easily transformable. In this study, we report a reduced transformation frequency of *B. subtilis* mutants lacking functional and structural flagellar components. This includes *hag*, the gene encoding the flagellin protein forming the filament of the flagellum. We confirm that the observed decrease of the transformation frequency is due to reduced expression of competence genes, particularly of the main competence regulator *comK*. The impaired competence is due to an increase in the phosphorylated form of the response regulator DegU, which is involved in regulation of both flagellar motility and competence. Altogether, our study identified a close link between motility and natural competence in *B. subtilis* suggesting that hindrance in motility has great impact on differentiation of this bacterium not restricted only to the transition towards sessile growth stage.

**Originality-Significance statement:** Understanding how versatile bacterial phenotypes influence each other is important for our basic understanding of microbial ecology. Our research highlights the novel intertwinement of bacterial differentiation and reveal how lack of single cell motility adjusts DNA exchange among bacterial strains.

## Introduction

When facing stressful environmental conditions, bacteria can respond with a variety of post-exponential modifications including secretion of degradative enzymes, sporulation, or genetic competence. *Bacillus subtilis* is one of the bacterial species that are able to take up free DNA from the environment and incorporate it into its own genome, a phenomenon referred to as natural competence (Dubnau, 1991). To import extracellular DNA into *B. subtilis* cells, a pseudopilus formed by proteins encoded by the *comG* operon facilitates binding to the receptor protein ComEA, which is located in the bacterial cell membrane (Inamine and Dubnau, 1995; Chen *et al.*, 2005). As only single stranded DNA is imported, the membrane-associated nuclease NucA catalyzes cleavage of the DNA after successful binding (Provvedi *et al.*, 2001). Subsequent transport of the DNA through a membrane channel formed by the protein ComEC is mediated by the ATPase ComFA that probably requires the transmembrane proton motive force (Maier *et al.*, 2004).

To take up DNA, cells have to be in a developmental state, in which a specific set of genes and regulators are expressed (Dubnau, 1991; Berka *et al.*, 2002). Regulation of the whole apparatus required for competence development is complex. Briefly, entry into the competence state occurs in a bistable manner during the early stationary phase, where a minority of cells produces high level of the competence master regulator ComK above a certain threshold that is required to switch on competence development, the so called ‘K-state’ (van Sinderen *et al.*, 1995; Maamar and Dubnau, 2005; Smits *et al.*, 2005; Dubnau and Losick, 2006). It was demonstrated that noise in the expression of *comK* determines the competent subpopulation and allows a dynamic stress response regarding competence development (Maamar *et al.*, 2007; Mugler *et al.*, 2016). Eventually, ComK activates the expression of late competence operons encoding the DNA-binding and-uptake machinery as well as genes, whose products are responsible for DNA integration (Berka *et al.*, 2002; Ogura *et al.*, 2002; Hamoen *et al.*, 2003). Increase of the ComK level is linked to a quorum sensing-mediated accumulation of the small ComS protein, which interferes with ComK degradation during the exponential phase (Turgay *et al.*, 1998). ComK is able to bind the *comK* promoter, triggering its own transcription, thus creating an auto-stimulatory loop (van Sinderen and Venema, 1994). This binding is further stabilized by the non-phosphorylated form of the regulator DegU to increase the level of ComK above the threshold sufficient for competence development (Hamoen *et al.*, 2000).

However, DegU is not only crucial for competence initiation, but also involved in the regulation of many other processes including protease production, biofilm development, and, particularly, flagellar motility (Murray *et al.*, 2009; Mukherjee and Kearns, 2014). The main components of the hook and basal body of the flagellum are encoded by the large *fla/che* operon. Transcription of this operon is activated by a complex formed by the regulator SwrA and phosphorylated DegU (DegU~P), which binds to one of the *fla/che* promoters (Mordini *et al.*, 2013; Mukherjee and Kearns, 2014). Amongst other genes, the operon contains the gene encoding the sigma factor σ^D^ that activates transcription of motility genes outside the *fla/che* operon like the *hag* gene (encoding flagellin), *motA* and *motB* (encoding flagellar stator proteins), as well as transcription of *lytF*, which is necessary for separation of motile cells after cell division (Serizawa *et al.*, 2004; Chen *et al.*, 2009). The level of σ^D^ and its position in the *fla*/*che* operon determines the cell fate, i.e. subpopulations of motile single cells or non-motile chains (Cozy and Kearns, 2010). The function of DegU~P changes in the absence of SwrA. In this case, DegU~P seems to inhibit motility via the same promoter of the *fla*/*che* operon (Amati *et al.*, 2004). Additionally, DegU~P can activate the anti-sigma factor FlgM by binding to its promoter region in the absence of SwrA (Hsueh *et al.*, 2011) allowing FlgM to antagonize σ^D^ (Caramori *et al.*, 1996). Consequently, DegU~P indirectly suppresses transcription of σ^D^-dependent genes (Hsueh *et al.*, 2011). It was suggested that a completion of flagellum assembly can be sensed by the DegSU two component system: FlgM, which is activated by DegU~P, causes inhibition of σ^D^-dependent genes, when the assembly of the flagellum is impeded (Cozy and Kearns, 2010; Hsueh *et al.*, 2011).

In addition to its role on modulating the expression of flagellum-related genes in *B. subtilis*, the phosphorylation and therefore the activity of DegU, has been shown to be influenced by a mechanical signal transmitted by the flagellum (Cairns *et al.*, 2013). Inhibition of flagellar rotation by the flagellar clutch or by tethering the flagella results in an increased DegU~P level in the cell.

In this study, we report a correlation between motility function and competence development, which in *B. subtilis* is connected by the multifunctional response regulator DegU. We show that mutants lacking a functional flagellum such as Δ*hag*, Δ*motA*, and Δ*flgE* exhibited a reduced transformation frequency. This was due to a decrease in competence gene expression, particularly reduced levels of the competence master regulator ComK, which can be reverted by overexpressing *comK* in the *hag* mutant. Finally, we suggest that the reduced transformation frequency was likely due to an imbalance in the phosphorylation level of DegU.

## Results

### Lack of active flagella impairs competence for DNA uptake in *B. subtilis*

While genetically modifying various *B. subtilis* strains, a striking difference in transformation frequency was observed between the wild type and a non-motile mutant lacking the gene encoding flagellin, *hag*. To explore this phenomenon, we tested the transformability of wild type (strain 168) and *hag*-mutant in competence medium (see Experimental Procedures), where the *hag*-mutant showed a more than 100-fold reduced transformation frequency relative to the wild type (Fig. 1A, B): while the transformation frequency of the wild type ranged between 3⋅10^−5^ and 5⋅10^−5^, that of the hag-mutant was reduced to values below 3⋅10^−7^. Similarly, the undomesticated *B. subtilis* strains DK1042 (transformable derivative of NCIB 3610) and PS216 showed reduced transformation efficiency when the *hag* gene of these strains was disrupted (Fig. S1). To investigate whether this difference in transformation frequency between the two strains resulted from a lower growth rate of the *hag*-mutant, the growth behavior of wild type and *hag*-mutant grown in competence medium was evaluated over time. As depicted in Fig. 1B, the *hag*-mutant showed a clear growth advantage and reached a higher OD compared to the wild type (unpaired two-sample t-test with Welch Correction: P = 0.001, n = 5), thus supporting our previous observations (Hölscher *et al.*, 2015). Further, it was tested whether the addition of DNA at different time points would increase the transformation frequency of the *hag*-mutant. However, the mutant showed a consistently low transformation frequency over the course of several hours, indicating that a shifted timing of the initiation of the competence state is unlikely to be the reason for the observed decrease in transformation frequency (Fig. 1C). To test whether this phenomenon is restricted to the *hag*-mutant or connected to the lack of an active motility apparatus in general, mutants lacking other functional flagellum-related genes were investigated. The transformation frequencies of mutants lacking the gene encoding one of the flagellar motor units, *motA*, and the gene encoding the hook protein, *flgE*, were decreased in both cases compared to the wild type (Fig. 2; unpaired two-sample t-test with Welch Correction: P = 0.01 for WT - Δ*motA*, P = 0.039 for WT - Δ*flgE*, n = 9 for both). Although the wild type transformation frequency was slightly different, the transformation frequencies of both Δ*motA* and Δ*flgE* were around 10-times lower than that of the wild type (Fig. 2). In contrast, a *cheA*-mutant lacking the main chemotaxis sensor kinase showed a similar transformation frequency than the wild type (Fig. S2; unpaired two-sample t-test with Welch Correction: P = 0.232, n = 3), suggesting that the presence of an active flagellum, but not directed motility *per se* is required for full competence development. In sum, these results demonstrate that the observed impaired competence is linked to a loss of flagellar function.

**Fig. 1.**
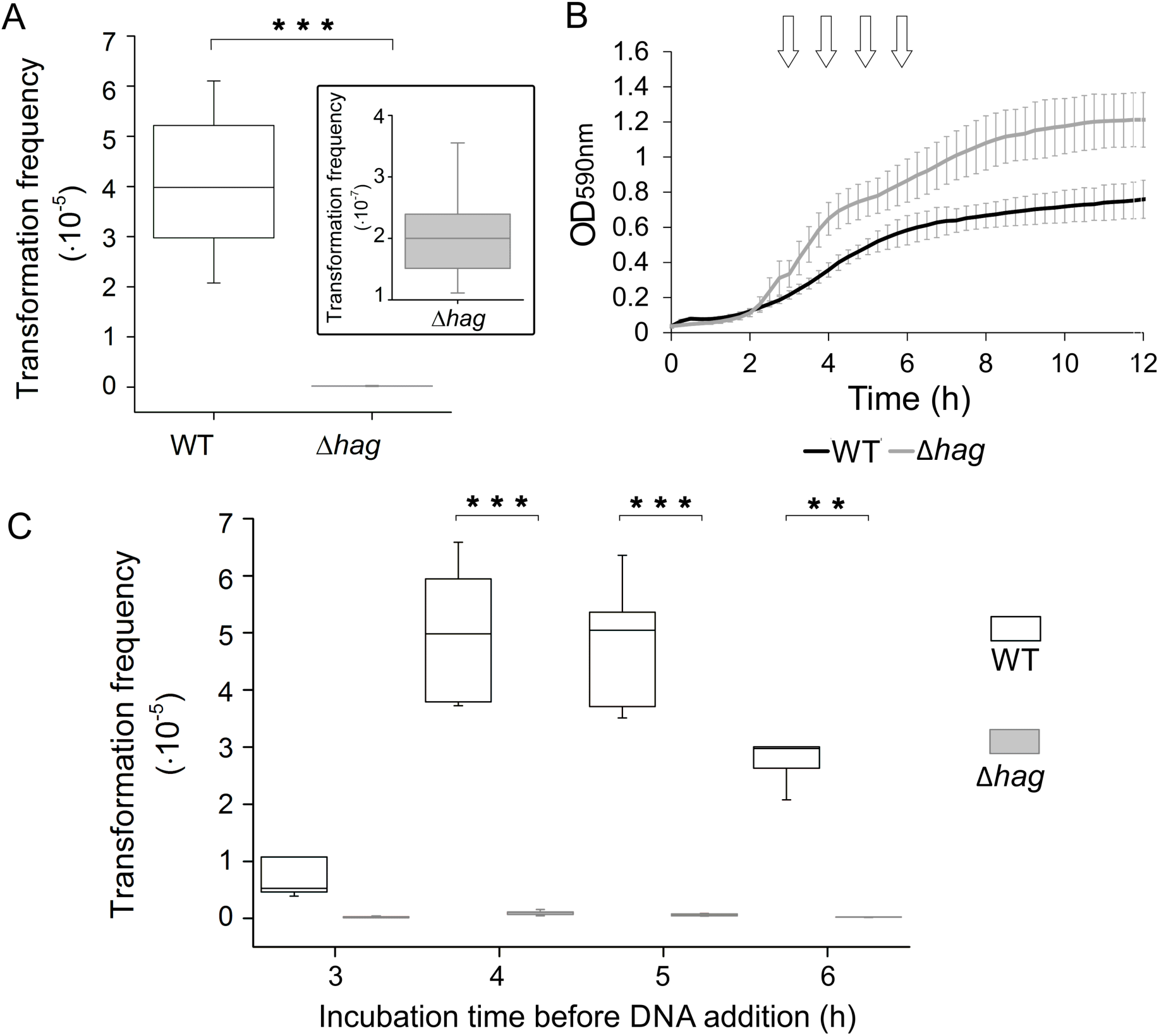
Transformation frequency is reduced in a mutant lacking flagellin protein. (A) Transformation frequency of *B. subtilis* wild type and *hag*-mutant after 6 h incubation in competence medium (unpaired two-sample t-test with Welch Correction: P = 3.1⋅10^−5^, n = 9). The inset shows a zoom-in of the *hag*-mutant data. (B) Growth dynamics of wild type and hag-mutant during 12 h incubation in competence medium. Standard deviations for the measurements are depicted in light grey (unpaired two-sample t-test with Welch Correction: P = 0.001, n = 5). Arrows indicate the time points of DNA addition to investigate the transformation frequency over time, which is shown as box-and-whisker plot in (C). The line in the boxes represents the median, the box indicates the 25^th^-75^th^ percentile. Asterisks indicate statistically significant differences between wild type and *hag*-mutant (unpaired two-sample t-test with Welch Correction for WT - Δ*hag*:, P = 0.125 for 3 h, P < 0.01 for 4 h, 5 h, 6 h; n = 6).

**Fig. 2.**
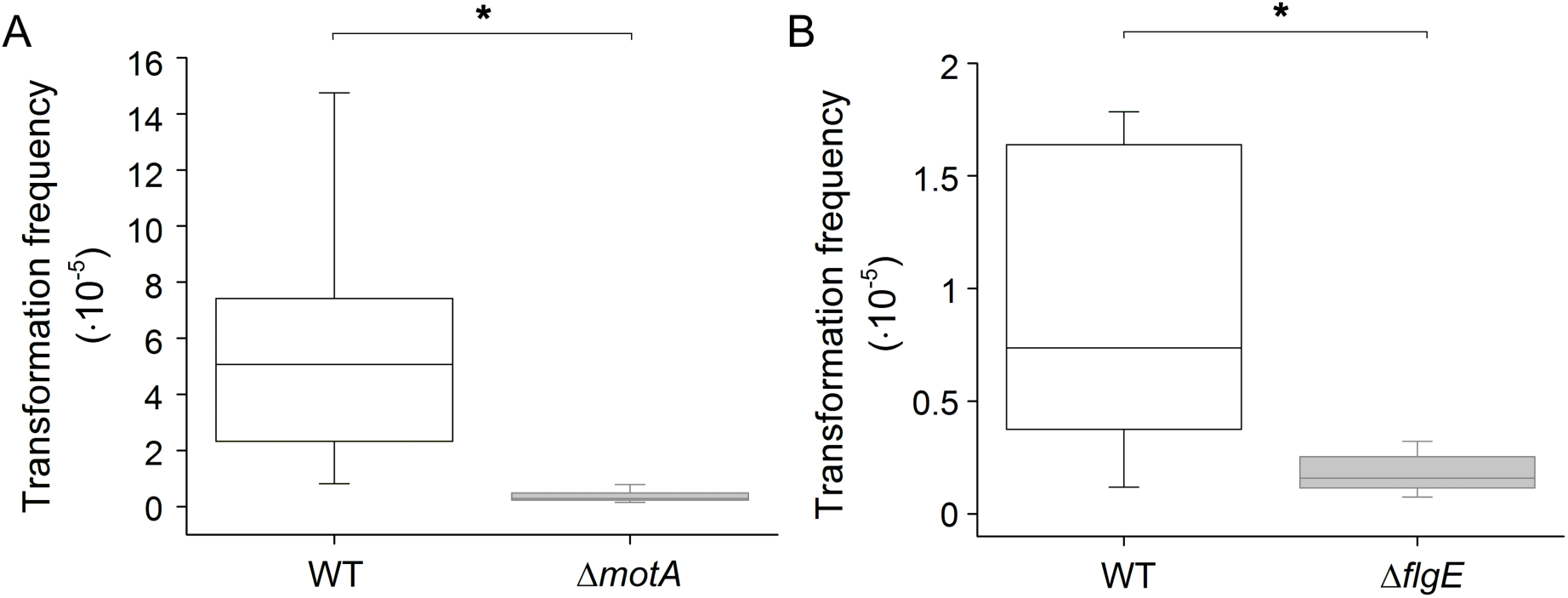
Mutants impaired in flagellar function exhibit lower transformation frequencies. Deletion of the gene encoding a flagellar stator (*motA*; A) or the gene encoding the hook protein (*flgE*; B) results in significantly lower transformation frequency of the respective strain compared to the wild type after incubation in competence medium for 6 h. The line in the boxes represents the median, the box indicates 25^th^-75^th^ percentile. Asterisks indicate statistically significant differences (unpaired two-sample t-test with Welch Correction: P = 0.01 for Δ*motA*; P = 0.039 for Δ*flgE*; n = 9 for all).

### Lack of competence in flagellar mutants is due to the reduced expression of competence genes

To determine if the detected diminished transformation frequency of flagellar mutants was due to altered competence gene expression, the fluorescent reporter P_*comG*_-*gfp* was introduced into these strains. This reporter allows the detection of cells expressing the *comG* operon-encoding genes required for pseudopilus formation and DNA uptake. In addition, this reporter provides a proxy on the activity of the ComK protein, the master regulator of competence. Qualitative microscopy analyses of cultures harboring the reporter, and which were grown in competence medium for 5 h, showed indeed a decreased number of fluorescent (i.e. *comG* expressing) cells in the *hag* mutant compared to the wild type, whereas a control strain lacking *comK* showed no fluorescence (Fig. 3A). For quantitative determination of competence gene expression within the population, flow cytometric measurements were performed that revealed 24.7% of fluorescent cells in wild type cultures (mean value), but only 4.6% of fluorescent cells for the *hag*-mutant (Fig. 3B, C; unpaired two-sample t-test with Welch Correction: P = 0.004, n = 3), thus confirming the microscopy results. Similarly, the *motA* and *flgE* mutants were analyzed microscopically as well as by using flow cytometry. Both methods revealed fewer cells activated transcription of competence genes in these mutants compared to the wild type (Fig. 4; unpaired two-sample t-test with Welch Correction: P = 0.017 for WT - Δ*motA*, P = 1.3⋅10^−9^ for WT - Δ*flgE*, n = 3 for both; mean percentage of fluorescent cells: 16.7% for wild type, 4.5% for Δ*motA*, 4.2% for Δ*flgE*). Flow cytometry measurements at different time points during growth in competence medium confirmed a similarly reduced fraction of competent cells in the *hag* mutant compared to the wild type strain (Fig. S3).

**Fig. 3.**
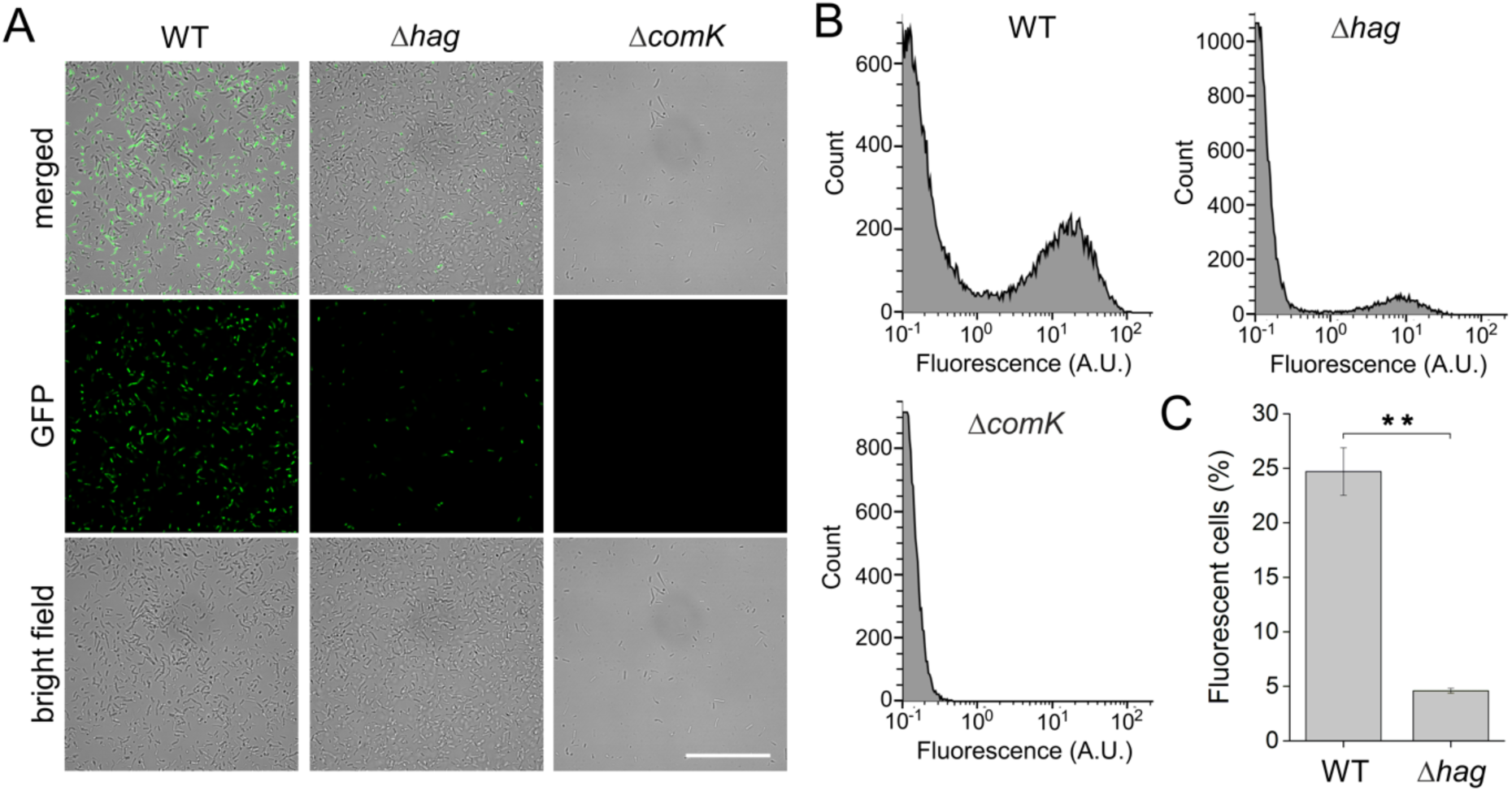
Fewer cells of the *hag*-mutant express competence genes compared to the wild type. (A) Representative microscopy images of strains harboring the P_*comG*_-gfp reporter in wild type, Δ*hag* or Δ*comK* genetic background. Images were recorded after incubation in competence medium for 5 h. The scale bar represents 50 µm. (B) Histograms of flow cytometric measurements showing the cell count and the fluorescence in arbitrary units for wild type, Δ*hag*, and Δ*comK* including background fluorescence. Representative images are shown for each strain. (C) Percentage of fluorescent cells determined from the data in (B) for wild type and *hag*-mutant by isolating the fluorescent population with fluorescence intensities above 3 A.U. Asterisks indicate significant differences (unpaired two-sample t-test with Welch Correction: P = 0.036, n = 3).

**Fig. 4.**
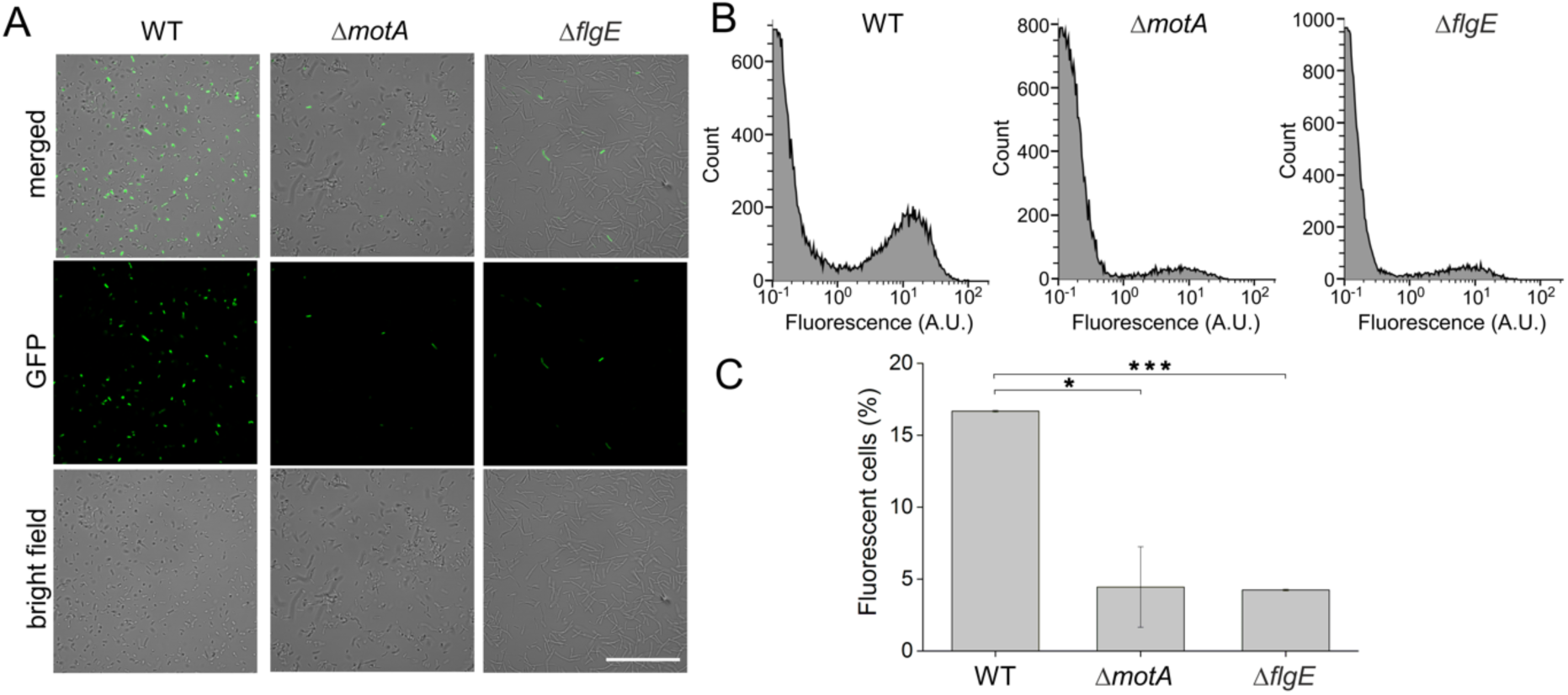
Competence gene expression is reduced in mutants lacking a functional flagellum. (A) Representative microscopy images of strains harboring the P_*comG*_-gfp reporter in wild type, Δ*motA*, or Δ*flgE* genetic background. Images were recorded after 5 h incubation in competence medium. The scale bar represents 50 µm. (B) Histograms of flow cytometric measurements showing the cell count and the fluorescence in arbitrary units for wild type, Δ*motA* or Δ*flgE*. Representative images are shown for each strain. (C) Percentage of fluorescent cells determined from the data in (B) by isolating the fluorescent population with fluorescence intensities above 3 A.U. showing a significant difference (asterisks) between wild type and Δ*motA* (P = 0.017) as well as wild type and Δ*flgE* (P < 0.001) with n = 3 for both (unpaired two-sample t-test with Welch Correction).

### Reduced competence in *hag* mutant can be rescued by overexpression of *comK*

The reduced competence gene expression in the tested flagellar mutants suggested a regulatory link between flagellar motility and competence. To investigate if regulatory elements upstream of *comK* were responsible for our observations and if a bypass of those could therefore rescue transformation frequency in the flagellar mutant, we examined a strain with an additional copy of *comK* under the control of a xylose-inducible promoter (P_*xyl*_-*comK*). Indeed, in combination with P_*xyl*_-*comK*, the transformation level of the *hag* mutant increased back to a level that was statistically indistinguishable from wild type levels (mean transformation frequency of 5.3⋅10^−6^ for the wild type and 8.5⋅10^−6^ for Δ*hag* P_*xyl*_-*comK*; Kruskal-Wallis test: P = 0.453, n = 9, Fig. 5A). Despite this observed increase in the *hag* strain upon *comK* overexpression, the wild type strain, which contained an inducible copy of *comK* showed a higher transformation frequency (Fig. 5A, Kruskal-Wallis test: P = 3.4⋅10^−4^ for WT – WT P_*xyl*_-*comK*, P = 3.4⋅10^−4^ for WT P_*xyl*_-*comK* - Δ*hag* P_*xyl*_-*comK*, n = 9 for both), which was probably due to higher levels of *comK* transcription at the native locus as previously observed (Hahn *et al*., 1996).

**Fig. 5.**
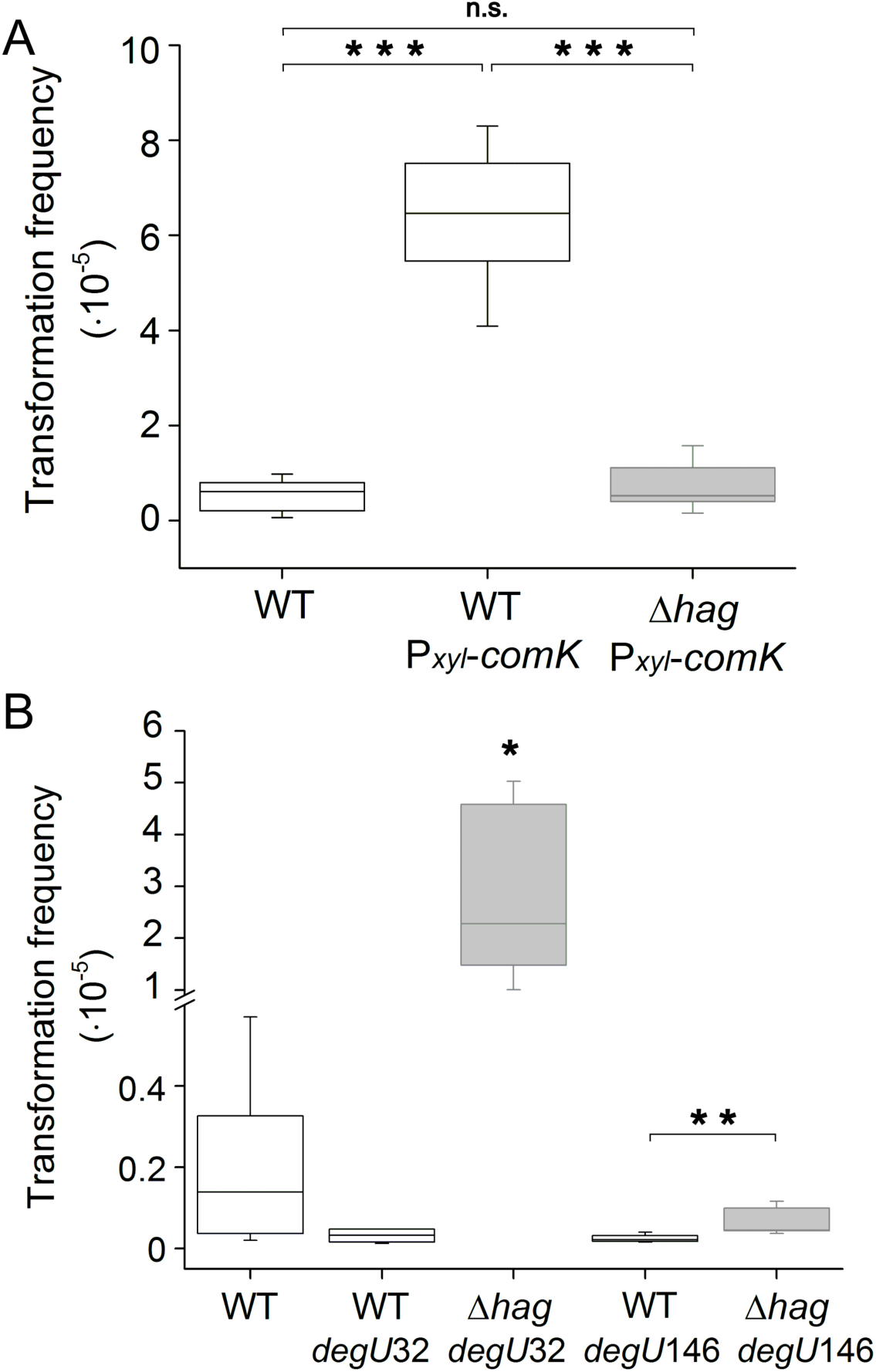
Synthetically induced *comK* and *degU*146 increase competence of Δ*hag*. (A) Transformation frequencies of wild type compared to strains harboring a xylose-inducible copy of *comK* (P_*xyl*_-*comK*) with wild type or Δ*hag* genetic background (Kruskal-Wallis test for WT - WT P_*xyl*_-comK: P = 3.4⋅10^−4^; for WT P_*xyl*_-*comK* - Δ*hag* P_*xyl*_-*comK*: P = 3.4⋅10^−4^, n = 9 for both). (B) Transformation frequencies of WT compared to strains harboring either a phosphorylated DegU variant (*degU*32) or a non-phosphorylatable DegU variant (*degU*146) in wild type or Δ*hag* background. Strain Δ*hag degU*32 is significantly different from all other strains (Kruskal-Wallis test: P < 0.05 for all, n=6). The line in the boxes represents the median, the box indicates 25^th^-75^th^ percentile. Asterisks indicate statistically significant differences (Kruskal-Wallis test: P = 0.007, n=6).

### Reduced competence in flagellar mutants is likely connected to unbalanced DegU phosphorylation

As the above results suggested that regulatory elements in response to impaired flagellar motility are responsible for the decreased *comK* expression, we investigated DegU as a likely candidate causing the reduced competence in flagellar mutants. As non-phosphorylated DegU was implicated to be required for *comK* transcription (Dahl *et al.*, 1992; Hamoen *et al.*, 2000), two variants of *degU* were tested: *degU*32, which harbors a mutation resulting in an extended half-life and thus higher stability of the phosphorylated form of the DegU protein (DegU~P), and *degU*146, which is cannot be phosphorylated (Dahl *et al.*, 1991; Dahl *et al.*, 1992; Kunst *et al.*, 1994). Both variants were tested in wild type as well as the Δ*hag* background to observe differences in transformability compared to the wild type strain. The results of this experiment indicated that the transformation frequency of the *degU*32 strain was slightly decreased (Figure 5B), which is consistent with previous publications, suggesting that non-phosphorylated DegU is required for priming *comK* transcription. The observed difference, however, was only marginally significant in our experimental setup (Figure 5B; Kruskal-Wallis test: P = 0.078, n = 6). Surprisingly, when combined with the Δ*hag* mutation, the transformability of *degU*32 increased significantly far above wild type levels, despite presumably possessing low levels of non-phosphorylated DegU to induce the ComK auto-stimulatory loop (Fig. 5B; Kruskal-Wallis test: P = 0.004, n = 6). Furthermore, we observed a tendency towards a reduced albeit non-significant transformation frequency in the *degU*146 strain compared to the wild type (Fig. 5B; Kruskal-Wallis test: P>0.05, n = 6). This result was similar to the one observed for *degU*32, although no negative impact on transformability was expected in strain *degU*146 due to the abolished phosphorylation of DegU. Interestingly, the *degU*146 strain combined with the Δ*hag* mutation exhibited transformation frequencies at the same level than the wild type strain (Fig. 5B; Kruskal-Wallis test: P>0.05, n = 6) that was significantly higher than the transformation frequency of the single *degU*146 mutant (Fig. 5B; Kruskal-Wallis test, P = 0.007, n = 6). These results suggest that altering the phosphorylation state of DegU in flagellar mutants can revert the negative impact on competence, which was caused by a lack of motility.

### Increased viscosity enhances competence in *B. subtilis*

A recent study showed that restricting the flagellar rotation by viscous medium results in induction of flagellar gene transcription and activation of the DegSU two-component system in *Paenibacillus* sp. NAIST15-1 (Kobayashi *et al.*, 2017). Accordingly, we tested whether an increased viscosity of the medium changes the transformability in *B. subtilis*. Indeed, the average transformation frequency of the wild type strain was three-fold higher in a medium of increased viscosity. The corresponding statistical test, however, indicated only a trend towards a statistically significant difference (Fig. 6; unpaired two-sample t-test: P = 0.095, n = 4).

**Fig. 6.**
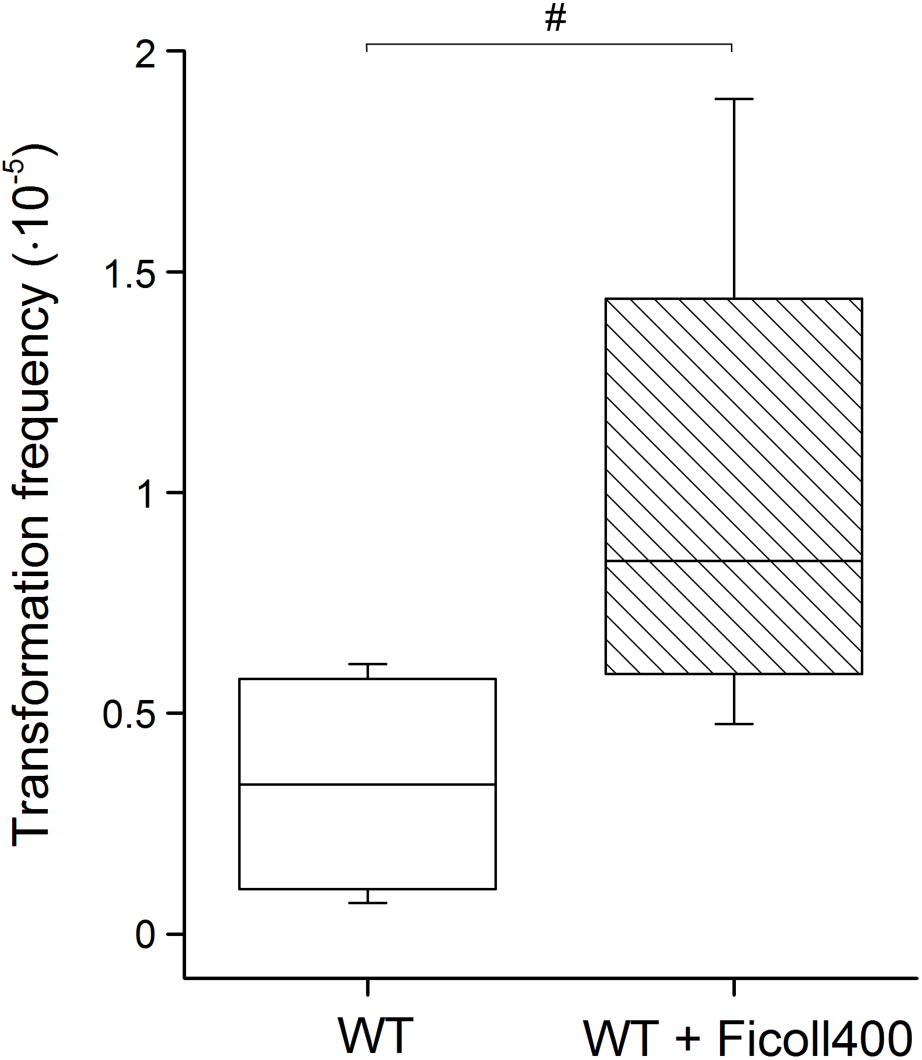
Competence is improved in viscous medium. Transformation frequency of the wild type strain grown in normal competence medium and in medium with increased viscosity. The line in the boxes represents the median, the box indicates 25^th^-75^th^ percentile, # indicates marginally significant differences (unpaired two-sample t-test: P = 0.095, n = 4).

## Discussion

Many cellular processes in *B. subtilis* are tightly connected through their underlying regulatory networks. Examples include motility and biofilm formation or biofilm formation and sporulation (e.g. Vlamakis *et al.*, 2013; Marlow *et al.*, 2014; Hölscher *et al.*, 2015). Here, we report an additional connection between flagellar motility and competence development. We could show that mutants with impaired flagellar function have defects in competence development. Such mutants displayed a considerably lower transformation frequency and expression of late competence genes, suggesting that it is due to an altered expression of the competence master regulator gene *comK*. The rescue experiment with an inducible *comK* confirmed that indeed competence could be rescued in the Δ*hag* strain, since Δ*hag* P_*xyl*_-*comK* exhibited a wild type transformation level.

In a recently published study, similar effects were observed even though different methods have been used: Diethmaier and colleagues found that the expression of *comK* is lower in deletion mutants of the *fla/che* operon, *hag*, and the second stator gene *motB* (Diethmaier *et al.*, 2017). While in their study the expression of *comK* was primarily monitored using a *comK* promoter fusion, our experiments predominantly assayed transformation frequency. Both studies, however, report a negative effect of the deletion of flagellar components on competence development. The extent to which wild type and mutant differ in competence development is in the same order of magnitude between the studies: for example Diethmaier *et al*. observe a 10-fold reduced number of *comG* expressing-cells in Δ*hag* (Diethmaier *et al.*, 2017), whereas our flow cytometry experiments showed a slightly lower, 5-fold reduction. Additionally, by investigating the transformation frequency in a *cheA* mutant, we could also show that the chemotactic response does not seem to have an influence on competence development.

Investigating modified variants of the response regulator DegU, we found that the transformation frequency of the *hag*-mutant could be restored to wild type level when the mutant carried a non-phosphorylatable DegU variant (*degU*146). This result suggests that a high level of DegU~P in the flagellar mutants was the reason for the decreased expression of the competence genes and *comK*, which could be counteracted by introducing a non-phosphorylatable variant of DegU. By additionally investigating a strain harboring a *degU-yfp* fusion, Diethmaier *et al*. also suggested an increased level of DegU~P to be present in the *hag*-mutant (Diethmaier *et al.*, 2017), which is consistent with our conclusions. In addition, the authors detected a reduced expression of *comK* in a strain with the *degU*32 variant, which produces a form of DegU~P with higher stability (Diethmaier *et al.*, 2017). Comparable results were obtained by Msadek *et al*., who found that high levels of DegU~P inhibit competence (Msadek *et al.*, 1990). We observed a similar, although weak statistical trend towards a reduced transformation frequency in *degU*32 strain. Miras and Dubnau (2016) have recently highlighted that differences in the DegU phosphorylation pathway among diverse *B. subtilis* isolates were likely responsible for variance in DNA transformation efficiency among certain domesticated and undomesticated strains. Moreover, slight differences in competence induction levels could also be affected by strain-specific characteristics. For example, *B. subtilis* 168 strains derived from different laboratories can exhibit striking variations in biofilm robustness (Gallegos-Monterrosa *et al.*, 2016). As suggested by Diethmaier and colleagues, the reduced transformation frequency in *degU*32 might be caused by the high DegU~P levels of this strain. However, the *degU*32 strain exhibits a non-motile phenotype and in the undomesticated strains DegU32 is not able to interact with SwrA at the P_A_ promoter of the *fla/che* operon (*swrA* is inactive in domesticated strains), leading to repression of P_A_ (*fla/che*) (Amati *et al.*, 2004; Mordini *et al.*, 2013). Due to low or no expression of the basic flagellar genes, this phenotype could mimic the situation observed in the flagellar mutants. In addition, we observed an increased transformation frequency when the *hag* gene was deleted from the *degU*32 background. This is in contrast to the model assuming that increased levels of phosphorylated DegU in the cells lowers competence. Therefore, it is possible that yet unidentified factors are also involved in connecting motility and competence development that might be independent of DegU~P. At this point however, we cannot provide a reasonable explanation for the increased transformation frequency of Δ*hag degU*32.

Interestingly, induction of competence state has negative impact on motility in *B. subtilis*. ComK negatively controls *hag* gene expression by stimulating the transcription of *comFA-C* operon and the downstream located anti-sigmaD factor coding gene, *flgM* (Liu and Zuber, 1998). This feedback loop presents another intriguing connection between these two cellular processes.

Diethmaier *et al*. proposed that increased DegU~P and lower *comK* expression in the flagellar mutants and in a strain with straight flagella was caused by a lower viscous load (Diethmaier *et al.*, 2017). In line with this report, we also observed that higher viscosity in the medium resulted in an increased transformation frequency. Nevertheless, a possible role of the DegSU two-component system in sensing incomplete assembly of flagella and dysfunction as suggested previously (Hsueh *et al.*, 2011; Cairns *et al.*, 2013) could also explain the increased DegU~P levels in the flagellar mutants.

Together, our results identify a connection between two major physiological processes, providing another example of the complexity of intracellular regulatory networks and the vast amount of tasks a single regulator can cover.

## Experimental Procedures

### Strains and cultivating conditions

The strains used in this study and their mutant derivatives are listed in Table S1. Mutants constructed in this study were obtained by natural transformation of a *B. subtilis* receptor strain with genomic DNA from a donor strain. Strain TB831 was created by transformation of strain 168 P_*xyl*_-*comK* with genomic DNA of strain GP902 (J. Stülke lab collection). To obtain strains TB926 and TB925, genomic DNA of strain 168 P_*comG*_-*gfp* was used to transform strain TB710 and TB689, respectively. Strain TB928 was obtained by transforming strain 168 P_*xyl*_-*comK* with genomic DNA of GP901 (J. Stülke lab collection). To create strain TB935 and TB936, strain 168 was transformed with genomic DNA obtained from strain QB4371 (Kunst *et al.*, 1994) and QB4458 (Dahl *et al.*, 1991), respectively. Their derivatives harbouring also a mutation of *hag* (TB923 and TB924) were created by transformation with genomic DNA, which was obtained from GP901. In-frame deletions of *motA*, *flgE*, and *cheA* were created using plasmids pEC1, pDP306, and pDP338, respectively, as previously described (Courtney *et al.*, 2012; Chan *et al.*, 2014; Calvo and Kearns, 2015). Strains were verified by fluorescence microscopy (P_*comG*_-*gfp* reporter), PCR (*hag* mutants), or sequencing (*degU* variants), using the oligonucleotides listed in Table S2. For experiments with strains harboring the inducible construct P_*xyl*_-*comK*, 1% of xylose (final concentration) was added for induction (see van den Esker *et al.*, 2017). To increase medium viscosity, 10% Ficoll400 (Carl Roth) was added to the medium before culture inoculation and the mix was vortexed vigorously for ca 20 s.

### Transformation frequency assay

To assess the transformation frequency of different strains, a modified version of the transformation protocol from Konkol *et al*. (2013) was used. 1 ml of each culture grown in 3 ml Lysogeny broth (LB) medium (LB-Lennox, Carl Roth; 10 g L^−1^ tryptone, 5 g L^−1^ yeast extract, and 5 g L^−1^ NaCl) for 16 h was centrifuged for 2 min at 11,000 x g. The pellet was washed twice in de-ionized water and was re-suspended in 100 µl de-ionized water. The re-suspended culture was diluted (1:80) in complete competence medium (MC: 1,8 ml de-ionized water, 6.7 µl 1M MgSO_4_, 50 µl 0.2% L-Tryptophan, 200 µl 10xMC; per 100 ml 10xMC: 14,036 g K_2_HPO_4_ [x3H_2_O], 5,239 g KH_2_PO_4_, 20 g glucose, 10 ml 300 mM tri-sodium citrate, 1 ml 83.97 mM ammonium iron (III) citrate, 1 g casein hydrolysate, 2 g potassium glutamate [H_2_O]) and incubated at 37°C, 225 rpm. For experiments with strains harboring P_*xyl*_-*comK*, 10x MC with fructose instead of glucose was used. After 6 h incubation time, 5 µl DNA with an antibiotic marker (PY79 *safA*::Tet gDNA, 60 ng/µl) was added to 500 µl culture. Any alteration in incubation time before addition of the DNA is indicated in the results section. Each culture was incubated for 30 min, then 500 µl fresh LB medium was added and the culture was incubated for another 1 h under the conditions mentioned above. Serial dilutions of cultures supplemented with DNA were prepared and plated on LB medium supplemented with 1.5 % agar to determine the number of colony forming units (cfu). Additionally, 50 µl and 100 µl undiluted cultures supplemented with gDNA as well as controls were plated on tetracycline (Tet) containing LB-agar plates (10 µg ml^−1^ Tet) to determine the number of transformant colonies. The transformation frequency was calculated by dividing the number of transformants per ml by cfu per ml.

### Growth curve experiments

To examine growth properties, cultures were inoculated in LB medium from frozen glycerol stocks and incubated for ca. 16 h at 37°C shaking at 225 rpm. Cultures were diluted 1:100 in 200 µl fresh completed MC medium (see above) and the OD_590nm_ was recorded for 16 h using a TECAN Infinite F200 PRO microplate reader. The cultures were incubated with orbital shaking with a duration of 800 s and an amplitude of 3 mm at 37°C and the OD_590_ was measured every 15 min.

### Fluorescence microscopy

Strains were investigated using a confocal laser scanning microscope (LSM 780, Carl Zeiss) equipped with an argon laser and a Plan-Apochromat/1.4 Oil DIC M27 63× objective. Cultures were grown prior microscopy for 5 h (if not indicated otherwise) in competence medium under the same conditions as described above (see section transformation frequency assay). Excitation of the fluorescent reporter (GFP) was performed at 488nm and the emitted fluorescence was recorded at 493-598nm. For image visualization, Zen 2012 software (Carl Zeiss) was used, brightness and contrast were adjusted equally in all images.

### Flow cytometry

Flow cytometric measurements were performed using a Partec CyFlow^®^ Space (Sysmex Partec GmbH, Germany), which was equipped with a solid-state laser for excitation of green/yellow fluorescent proteins at 488 nm. Single cells were detected in forward and sideward scatter channels as well as in one fluorescent channel. A minimum of 40,000 cells were analyzed for the experiments. To define the background fluorescence signal, non-labelled *B. subtilis* cultures were analyzed as control. Cultures used for measurements were grown for 5 h (if not indicated otherwise) in competence medium under the same conditions as described above (see section transformation frequency assay). For evaluation of the data, the FlowJo® software (FlowJo LLC, Ashland, USA) was used and a gate was set at 3 fluorescence units for all samples to isolate the fluorescent population and determine the percentage of fluorescent cells.

### Statistics

Statistical analyses were performed using OriginPro 2016 (V93E, OriginLab Northampton, USA). Unpaired two-sample t-test with Welch Correction or a Kruskal-Wallis test was used to test for significant differences.

## Acknowledgement

We thank Daniel Kearns for providing plasmids, Anne Richter for constructing the in-frame deletion mutants, and Tadeja Marin for initial observations. We thank Wiep Klaas Smits for his comment on our bioRxiv submission. This project was funded by grant KO4741/3-1 from the German Research Foundation (DFG). T.H. and A.D. were supported by International Max Planck Research School and Alexander von Humboldt foundation fellowships, respectively. CK was supported by the Volkswagen Foundation (I/85 290) as well as the German Research Foundation (SFB 944/2-2016, KO 3909/2-1). Á.T.K. was supported by a Startup fund from the Technical University of Denmark.

## References

Amati, G., Bisicchia, P., and Galizzi, A. (2004) DegU-P represses expression of the motility *fla-che* operon in *Bacillus subtilis*. J Bacteriol 186: 6003–6014.

Berka, R.M., Hahn, J., Albano, M., Draskovic, I., Persuh, M., Cui, X., et al. (2002) Microarray analysis of the Bacillus subtilis K-state: Genome-wide expression changes dependent on ComK. Mol Microbiol 43: 1331–1345.

Cairns, L.S., Marlow, V.L., Bissett, E., Ostrowski, A., and Stanley-Wall, N.R. (2013) A mechanical signal transmitted by the flagellum controls signalling in Bacillus subtilis. Mol Microbiol 90: 6–21.

Calvo, R.A., and Kearns, D.B. (2015) FlgM is secreted by the flagellar export apparatus in Bacillus subtilis. J Bacteriol 197: 81–91.

Caramori, T., Barilla, D., Nessi, C., Sacchi, L., and Galizzi, A. (1996) Role of FlgM in sigma(D)-dependent gene expression in Bacillus subtilis. J Bacteriol 178: 3113–3118.

Chan, J.M., Guttenplan, S.B., and Kearns, D.B. (2014) Defects in the flagellar motor increase synthesis of poly-γ-glutamate in Bacillus subtilis. J Bacteriol 196: 740–753.

Chen, I., Christie, P.J., and Dubnau, D. (2005) The ins and outs of DNA transfer in bacteria. Science 310: 1456–1460.

Chen, R., Guttenplan, S.B., Blair, K.M., and Kearns, D.B. (2009) Role of the σD-dependent autolysins in Bacillus subtilis population heterogeneity. J Bacteriol 191: 5775–5784.

Courtney, C.R., Cozy, L.M., and Kearns, D.B. (2012) Molecular characterization of the flagellar hook in Bacillus subtilis. J Bacteriol 194: 4619–4629.

Cozy, L.M., and Kearns, D.B. (2010) Gene position in a long operon governs motility development in Bacillus subtilis. Mol Microbiol 76: 273–285.

Dahl, M.K., Msadek, T., Kunst, F., and Rapoport, G. (1991) Mutational analysis of the Bacillus subtilis DegU regulator and its phosphorylation by the DegS protein kinase. J Bacteriol 173: 2539–2547.

Dahl, M.K., Msadek, T., Kunst, F., and Rapoport, G. (1992) The phosphorylation state of the DegU response regulator acts as a molecular switch allowing either degradative enzyme synthesis or expression of genetic competence in Bacillus subtilis. J Biol Chem 267: 14509–14514.

Diethmaier, C., Chawla, R., Canzoneri, A., Kearns, D.B., and Dubnau, D. (2017) Viscous drag on the flagellum activates Bacillus subtilis entry into the. Mol Microbiol 106: 367–380.

Dubnau, D. (1991) Genetic competence in Bacillus subtilis. Microbiol Rev 55: 395–424.

Dubnau, D., and Losick, R. (2006) Bistability in bacteria. Mol Microbiol 61: 564–572.

van den Esker, M.H., Kovács, Á.T., and Kuipers, O.P. (2017) YsbA and LytST are essential for pyruvate utilization in Bacillus subtilis. Environ Microbiol 19: 83–94.

Gallegos-Monterrosa, R., Mhatre, E., and Kovács, Á.T. (2016) Specific Bacillus subtilis 168 variants form biofilms on nutrient-rich medium. Microbiol 162: 1922–1932.

Hamoen, L.W., Venema, G., and Kuipers, O.P. (2003) Controlling competence in Bacillus subtilis: shared use of regulators. Microbiol 149: 9–17.

Hamoen, L.W., Werkhoven, a F. Van, Venema, G., and Dubnau, D. (2000) The pleiotropic response regulator DegU functions as a priming protein in competence development in Bacillus subtilis. Proc Natl Acad Sci U S A 97: 9246–9251.

Hölscher, T., Bartels, B., Lin, Y.-C., Gallegos-Monterrosa, R., Price-Whelan, A., Kolter, R., et al. (2015) Motility, chemotaxis and aerotaxis contribute to competitiveness during bacterial pellicle biofilm development. J Mol Biol 427: 3695–3708.

Hsueh, Y.H., Cozy, L.M., Sham, L.T., Calvo, R.A., Gutu, A.D., Winkler, M.E., and Kearns, D.B. (2011) DegU-phosphate activates expression of the anti-sigma factor FlgM in Bacillus subtilis. Mol Microbiol 81: 1092–1108.

Inamine, G.S., and Dubnau, D. (1995) ComEA, a Bacillus subtilis integral membrane protein required for genetic transformation, is needed for both DNA binding and transport. J Bacteriol 177: 3045–3051.

Kobayashi, K., Kanesaki, Y., and Yoshikawa, H. (2017) Surface sensing for flagellar gene expression on solid media in Paenibacillus sp. NAIST15-1. Appl Environ Microbiol 83: e00585–17.

Konkol, M.A., Blair, K.M., and Kearns, D.B. (2013) Plasmid-encoded comi inhibits competence in the ancestral 3610 strain of Bacillus subtilis. J Bacteriol 195: 4085–4093.

Kunst, F., Msadek, T., Bignon, J., and Rapoport, G. (1994) The DegS/DegU and ComP/ComA two-component systems are part of a network controlling degradative enzyme synthesis and competence in Bacillus subtilis. Res Microbiol 145: 393–402.

Liu, J., and Zuber, P. (1998) A molecular switch controlling competence and motility: competence regulatory factors ComS, MecA, and ComK control sigmaD-dependent gene expression in Bacillus subtilis. J Bacteriol 180: 4243–4251.

Maamar, H., and Dubnau, D. (2005) Bistability in the Bacillus subtilis K-state (competence) system requires a positive feedback loop. Mol Microbiol 56: 615–24.

Maamar, H., Raj, A., and Dubnau, D. (2007) Noise in Gene Expression Determines. Science 526–529.

Maier, B., Chen, I., Dubnau, D., and Sheetz, M.P. (2004) DNA transport into Bacillus subtilis requires proton motive force to generate large molecular forces. Nat Struct Mol Biol 11: 643–649.

Marlow, V.L., Porter, M., Hobley, L., Kiley, T.B., Swedlow, J.R., Davidson, F. a, and Stanley-Wall, N.R. (2014) Phosphorylated DegU manipulates cell fate differentiation in the Bacillus subtilis biofilm. J Bacteriol 196: 16–27.

Miras, M., and Dubnau, D. (2016) A DegU-P and DegQ-Dependent Regulatory Pathway for the K-state in Bacillus subtilis. Front Microbiol 7: 1–14.

Mordini, S., Osera, C., Marini, S., Scavone, F., Bellazzi, R., Galizzi, A., and Calvio, C. (2013) The Role of SwrA, DegU and PD3 in fla/che Expression in B. subtilis. PLoS One 8: e85065.

Msadek, T., Kunst, F., Henner, D., Klier, A., Rapoport, G., and Dedonder, R. (1990) Signal transduction pathway controlling synthesis of a class of degradative enzymes in Bacillus subtilis: expression of the regulatory genes and analysis of mutations in degS and degU. J Bacteriol 172: 824–834.

Mugler, A., Kittisopikul, M., Hayden, L., Liu, J., Wiggins, C.H., Süel, G.M., and Walczak, A.M. (2016) Noise expands the response range of the Bacillus subtilis competence circuit. PLoS Comput Biol 12: 1–21.

Mukherjee, S., and Kearns, D.B. (2014) The structure and regulation of flagella in Bacillus subtilis. Annu Rev Genet 48: 319–40.

Murray, E.J., Kiley, T.B., and Stanley-Wall, N.R. (2009) A pivotal role for the response regulator DegU in controlling multicellular behaviour. Microbiol 155: 1–8.

Ogura, M., Yamaguchi, H., Kobayashi, K., Ogasawara, N., and Fujita, Y. (2002) Whole-genome analysis of genes regulated by the Bacillus subtilis competence transcription factor ComK. J Appl Microbiol 184: 2344–2351.

Provvedi, R., Chen, I., and Dubnau, D. (2001) NucA is required for DNA cleavage during transformation of Bacillus subtilis. Mol Microbiol 40: 634–644.

Serizawa, M., Yamamoto, H., Yamaguchi, H., Fujita, Y., Kobayashi, K., Ogasawara, N., and Sekiguchi, J. (2004) Systematic analysis of SigD-regulated genes in Bacillus subtilis by DNA microarray and Northern blotting analyses. Gene 329: 125–136.

van Sinderen, D., Luttinger, A., Kong, L., Dubnau, D., Venema, G., and Hamoen, L. (1995) comK encodes the competence transcription factor, the key regulatory protein for competence development in Bacillus subtilis. Mol Microbiol 15: 455–462.

van Sinderen, D., and Venema, G. (1994) ComK acts as an autoregulatory control switch in the signal transduction route to competence in Bacillus subtilis. J Bacteriol 176: 5762–5770.

Smits, W.K., Eschevins, C.C., Susanna, K.A., Bron, S., Kuipers, O.P., and Hamoen, L.W. (2005) Stripping Bacillus: ComK auto-stimulation is responsible for the bistable response in competence development. Mol Microbiol 56: 604–614.

Turgay, K., Hahn, J., Burghoorn, J., and Dubnau, D. (1998) Competence in Bacillus subtilis is controlled by regulated proteolysis of a transcription factor. EMBO J 17: 6730–6738.

Vlamakis, H., Chai, Y., Beauregard, P., Losick, R., and Kolter, R. (2013) Sticking together: building a biofilm the Bacillus subtilis way. Nat Rev Microbiol 11: 157–68.

